# The role of ATP hydrolysis in conformational changes of human P-glycoprotein in living cells

**DOI:** 10.1101/641225

**Authors:** Ryota Futamata, Fumihiko Ogasawara, Takafumi Ichikawa, Atsushi Kodan, Yasuhisa Kimura, Noriyuki Kioka, Kazumitsu Ueda

## Abstract

P-glycoprotein (P-gp; also known as MDR1 or ABCB1) is an ATP-driven multidrug transporter that extrudes various hydrophobic toxic compounds to the extracellular space. P-gp consists of two transmembrane domains (TMDs) that form the substrate translocation pathway and two nucleotide-binding domains (NBDs) that bind and hydrolyze ATP. P-gp takes at least two states during transport; the inward-facing (pre-drug transport) conformation, in which the two NBDs are separated and the two TMDs are open to the intracellular side, and the outward-facing (post-drug transport) conformation, in which the NBDs are dimerized and the TMDs are slightly open to the extracellular side. ATP binding and hydrolysis cause conformational changes between the inward-facing and the outward-facing conformations to translocate substrates across the membrane. However, it remains unclear how ATP is used during these conformational changes in living cells. In this study, we investigated the role of ATP binding and hydrolysis during the conformational changes of human P-gp in living cells by using fluorescence resonance energy transfer (FRET). We show that ATP binding causes the conformational change to the outward-facing state and that ATP hydrolysis and subsequent release of γ-phosphate from both NBDs allow the outward-facing state to return to the original inward-facing state.

## Introduction

P-gp, a member of the ATP-binding cassette (ABC) transporter family, is an ATP-dependent efflux pump that transports various hydrophobic compounds (1, 2). Its substrates include therapeutic drugs, alkaloids, flavonoids, and other hydrophobic natural toxic compounds. Accordingly, P-gp is an essential component of the protective physiological barriers in important organs (3), such as the brain and testis. The protein consists of four core domains: two transmembrane domains (TMDs), which create the translocation pathway for the substrates, and two nucleotide-binding domains (NBDs), which bind and hydrolyze ATP to power the transport process. Although numerous studies have been conducted (4–8) following the discovery of P-gp (9–11), it remains unclear how ATP is utilized by P-gp to export hydrophobic compounds from cells.

Recently, we reported a pair of structures of the P-gp orthologue (CmABCB1) from the thermophilic unicellular eukaryote *Cyanidioschyzon merolae* at resolutions of 2.4 Å and 1.9 Å, respectively: an inward-facing apo state (12) and an outward-facing state with a bound nucleotide (13). Conformational changes from the inward-facing to the outward-facing state during drug transport could be clearly visualized at these high resolutions. In the inward-facing state, a large inner cavity was formed in the center of the TMDs, extending from the middle of the lipid bilayer to the cytosol, and the two NBDs were separated by about 26 Å. In the outward-facing state, the NBDs were in close proximity, resulting in the formation of nucleotide binding sites composed of a Walker A motif (P-loop) in one NBD and a signature motif in the other NBD. Conformational changes in the NBDs caused by nucleotide binding mediated TMD movements. The outward-facing structure at 1.9 Å resolution (13) revealed that a relay of van der Waals interactions and hydrogen-bond networks formed by ATP binding propagate from the NBDs to the TMDs and make the whole molecule move as a rigid body. On the other hand, the inward-facing structure (12) contains two layers of networks, a hydrogen bond network and an aromatic hydrophobic network, at the top of the molecule, which stabilize the inward-facing state, but the networks between the NBDs and TMDs were not observed, which is unlike the outward-facing conformation. These structural features suggest that ATP binding in both NBDs drives conformational movements from the flexible inward-facing conformation (where the networks between the NBDs and TMDs are not formed) to the rigid outward-facing conformation (where the networks between the NBDs and TMDs are formed) of P-gp, and that ATP hydrolysis releases the rigid outward-facing conformation to the inward-facing conformation. This mechanism, in which ATP hydrolysis triggers conformational change from the outward-facing to the inward-facing state, is supported by a study using mouse P-gp reconstituted in lipid nanodiscs composed of a scaffold protein and lipid bilayer (7). However, few studies have been conducted to investigate the roles of ATP binding and ATP hydrolysis in the conformational changes of P-gp under a physiological membrane environment. In this study, we analyzed the roles of ATP binding and ATP hydrolysis in the conformational changes of human P-gp in living cells by using fluorescence resonance energy transfer (FRET).

## Results

### A FRET construct to detect the conformational change of human P-gp in living cells

The distances between the C-terminal of NBD1 and that of NBD2 of human P-gp are estimated to be about 30 Å and 11 Å in the inward-facing and outward-facing structures, respectively (Movie_S01). Because this difference in distance is assumed to be one of the largest between the inward-facing and outward-facing structures, we considered it suitable for FRET analysis in order to detect the conformational change of human P-gp in living cells in real time. We generated a FRET construct, P-gp–FRET, in which Cerulean (donor) was inserted after NBD1 and Venus (acceptor) was fused after NBD2 of human P-gp (Fig. 1A). Another FRET construct, P-gp–VsCn, in which Venus and Cerulean were fused tandemly after NBD2, was predicted to show a high level of FRET despite the conformation. P-gp–Cerulean, in which Cerulean was inserted after NBD1, and P-gp–Venus, in which Venus was fused after NBD2, were constructed as negative controls. When these constructs were transiently expressed in HEK293 cells, fluorescence signals were observed on the plasma membrane (Fig. S1A). Basal and strong signals were observed in the FRET channel with P-gp–FRET and P-gp–VsCn, respectively, whereas almost no signal was observed with P-gp–Cerulean or P-gp–Venus (Fig. S1A, right). When cells expressing these P-gp constructs were incubated with rhodamine 6G (R6G), a transport substrate of P-gp, the R6G accumulation was at a lower level than in mock-transfected cells but was restored by co-incubation with PSC-833, a specific inhibitor of P-gp (Fig. S1B). These results indicated that neither the insertion nor fusion of fluorescent proteins impaired the plasma membrane localization or function of P-gp. Due to the leakage of Venus fluorescence in the detection channel, the fluorescence of cells expressing P-gp constructs containing Venus were shifted to the right.

**Figure 1.**
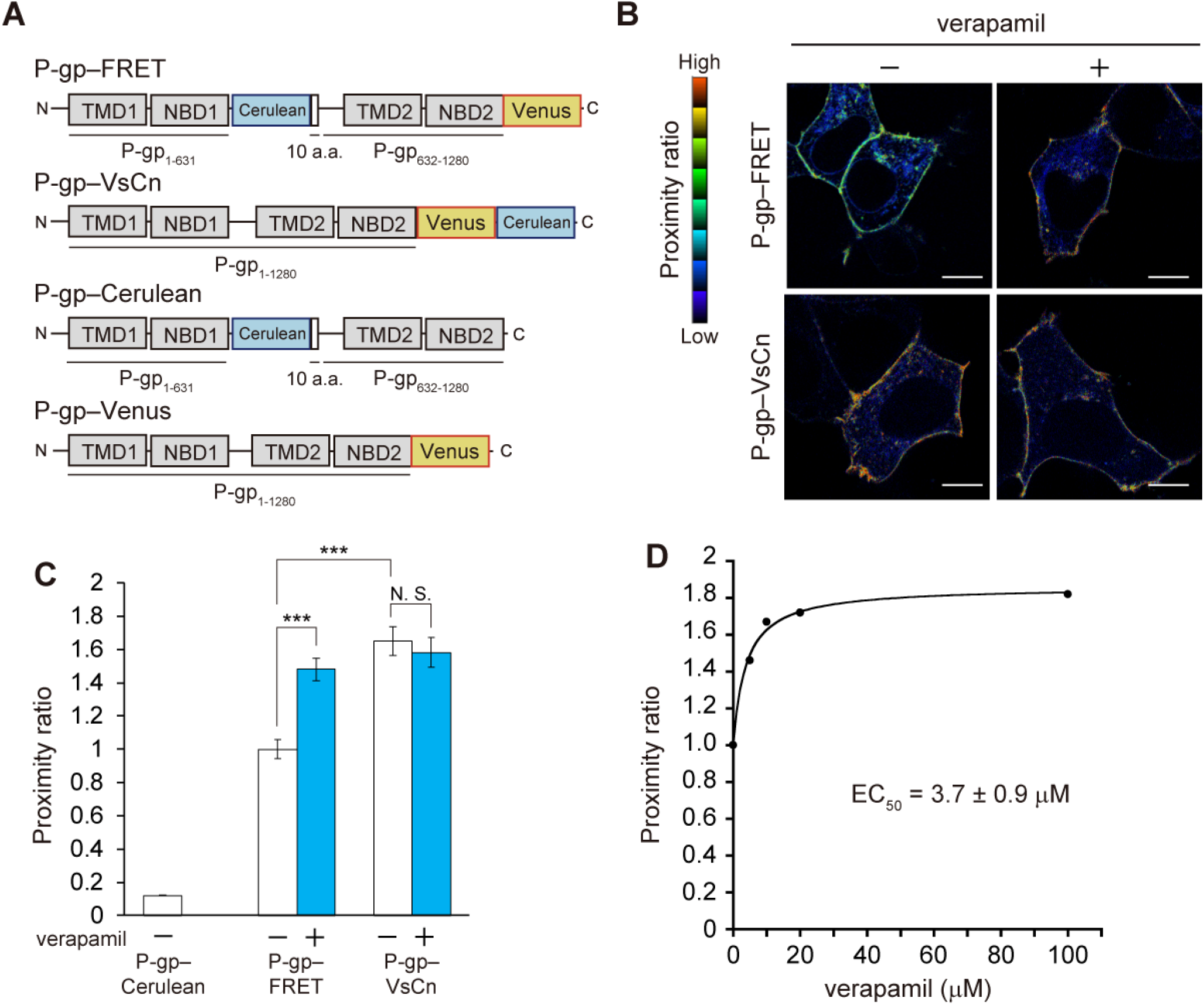
Effects of verapamil on the conformational changes of P-gp. (**A**) Schematic representations of the fluorescent protein–tagged P-gps. (**B**) Proximity ratio pseudocolor images of HEK293 cells expressing fluorescent protein–tagged P-gps. Cells were treated with DMSO or 100 μM verapamil for 5 min at 37°C. The proximity ratio is shown in an 8-color scale, and each hue has a range of 32 intensities based on the corrected intensity of the FRET image. Scale bar; 10 μm. (**C**) Quantification of the proximity ratio. The area corresponding to the plasma membrane was determined from automatically segmented Venus or Cerulean images (Fig. S3A), and then the proximity ratio on the plasma membrane was calculated. (**D**) Dose-dependent increase of the proximity ratio by verapamil. The proximity ratio is shown relative to that of P-gp–FRET. Data are expressed as means ± S.E.; n = 10–15 cells per sample; ***P < 0.001; N.S., P > 0.05.

### A transport substrate causes conformational changes in P-gp

Next, we investigated whether the addition of a transport substrate could affect the level of FRET. We defined the proximity ratio as the ratio of the corrected intensity of the FRET signal divided by the Cerulean intensity, which is shown in pseudocolor in Fig. 1B. The proximity ratio of P-gp–FRET on the plasma membrane (shown in green) was low, and the signal of P-gp–VsCn (shown in yellow or red) was high in the absence of a transport substrate. When verapamil, a transport substrate of P-gp (14) that stimulates its ATPase activity (Fig. S2), was added to the cells, the proximity ratio of P-gp–FRET increased, whereas that of P-gp–VsCn was unchanged (Fig. 1B). Next, the plasma membrane was extracted by automatic membrane-segmentation of the Venus image of P-gp–FRET, –VsCn or the Cerulean image of P-gp–Cerulean (Fig. S3A), and the proximity ratio on the plasma membrane was calculated. The proximity ratio of P-gp–FRET increased about 1.5-fold upon the addition of 100 μM verapamil, which approximates the level of P-gp–VsCn (Fig. 1C). Furthermore, the proximity ratio was not affected by the expression level of P-gp (Fig. S3B), suggesting that the increase upon verapamil treatment is independent of the intermolecular FRET caused by condensation of the FRET constructs. Verapamil increased the proximity ratio in a concentration-dependent manner, and the EC_50_ was 3.7 ± 0.9 μM (Fig. 1D). On the other hand, the proximity ratio of P-gp–VsCn was unaffected by verapamil, and that of P-gp–Cerulean was less than one-tenth that of P-gp–VsCn. These results suggest that verapamil causes conformational changes in P-gp.

### ATP binding- and hydrolysis-deficient mutants are trapped in low- and high-FRET states, respectively

To investigate the involvement of ATP binding and ATP hydrolysis in the conformational changes induced by verapamil, we constructed P-gp(MM)–FRET (Fig. 4E), in which conserved lysine residues in the Walker A motif crucial for ATP binding (8,13,15) were replaced by methionines, and P-gp(QQ)–FRET(Fig. 4F), in which glutamate residues in the Walker B motif crucial for ATP hydrolysis (13, 16) were replaced by glutamines. Both P-gp(MM)–FRET and P-gp(QQ)–FRET properly localized to the plasma membrane (Fig. S4A) but had no R6G transport activity (Fig. S4B). The proximity ratio of P-gp(MM)–FRET was comparable to that of P-gp–FRET in the absence of a transport substrate but was unaffected by verapamil treatment (Fig. 2A). This result suggests that ATP binding is required for the increase of the proximity ratio induced by verapamil. On the other hand, the proximity ratio of P-gp(QQ)–FRET, which allows ATP binding but reduces the rate of ATP hydrolysis (16), was significantly higher than that of P-gp–FRET even in the absence of a transport substrate and was also unaffected by verapamil (Fig. 2B). This result suggests that ATP hydrolysis causes a transition from the high FRET to the low FRET state. Like P-gp–VsCn, control constructs P-gp(MM)–VsCn and P-gp(QQ)–VsCn exhibited a high proximity ratio independent of verapamil.

**Figure 2.**
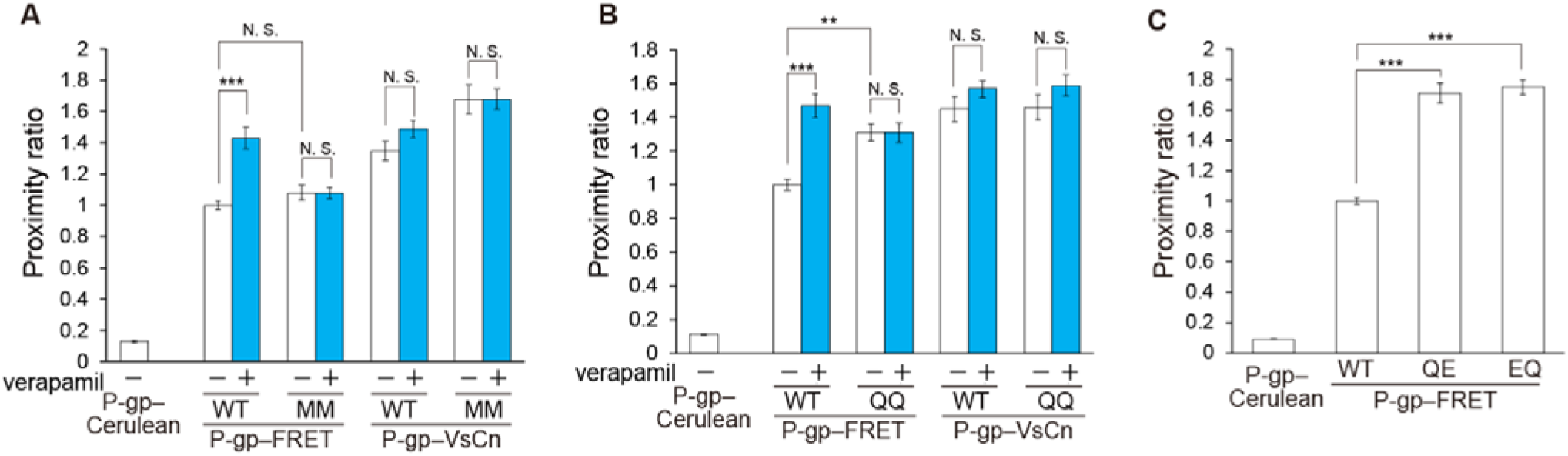
Proximity ratio of ATP binding-deficient and ATP hydrolysis-deficient mutants. The proximity ratio in intact cells of the ATP binding-deficient mutant MM (**A**) and ATP hydrolysis-deficient mutant QQ (**B**), and two single Walker B mutants, QE and EQ (**C**). HEK293 cells transiently transfected with fluorescent protein–tagged P-gps were treated with DMSO (empty) or 100 μM verapamil (blue) for 5 min at 37°C. The proximity ratios are normalized to that of P-gp–FRET. Data are expressed as means ± S.E.; n = 12–15 cells per sample; **P < 0.01; ***P < 0.001; N.S., P > 0.05.

**Figure 3.**
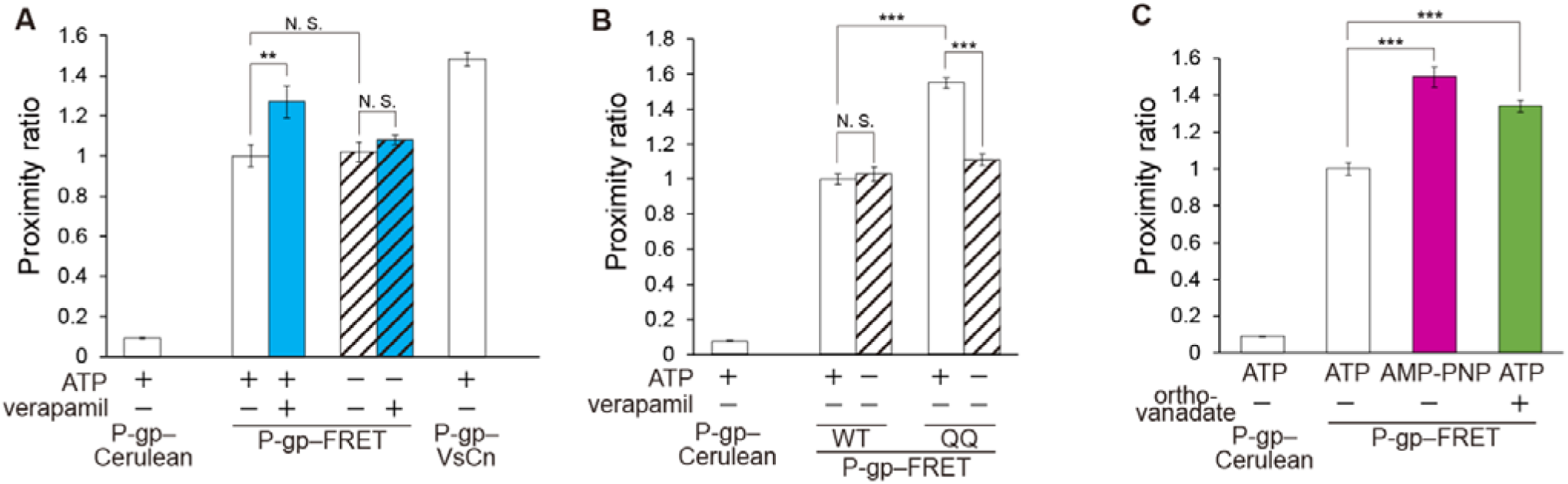
Effects of ATP binding and ATP hydrolysis on the conformational changes of P-gp. (**A**, **B**) The proximity ratio in permeabilized semi-intact cells treated with DMSO (empty) or 100 μM verapamil (blue) in the presence or absence of 4.5 mM MgATP (hatched). (**C**) The proximity ratio in permeabilized semi-intact cells in the presence of 4.5 mM MgATP (empty), 4.5 mM MgAMP-PNP (magenta), or 4.5 mM MgATP and 1 mM orthovanadate (green). The proximity ratios are normalized to that of P-gp–FRET. Data are expressed as means ± S.E.; n = 12–15 cells per sample; **P < 0.01; ***P < 0.001; N.S., P > 0.05.

### ATP binding is required for the conformational change of P-gp

To further demonstrate the role of ATP binding in the conformational change of P-gp, intracellular ATP was depleted from cells by treating them with streptococcal toxin streptolysin O (SLO). SLO forms 25–30 nm aqueous pores within the plasma membrane, which allows the free passage of ions and small molecules (17). 4’,6-diamidino-2-phenylindole (DAPI) hardly goes through the plasma membrane of intact cells. Therefore, cells with a DAPI-positive nucleus and permeabilized plasma membranes were used for the FRET analysis (Fig. S5). DAPI is not a transport substrate of P-gp and does not stimulate its ATPase activity (Fig. S2). In the presence of exogenously added ATP, the proximity ratio of P-gp–FRET was increased by the addition of verapamil (Fig. 3A), consistent with the observations for intact cells (Fig. 1C). However, in the absence of ATP, the proximity ratio of P-gp–FRET was not increased even after the addition of verapamil. These results suggest that both ATP and substrate binding are required for enhancing the conformational change from the inward-facing state (Fig. 4A-D).

**Figure 4.**
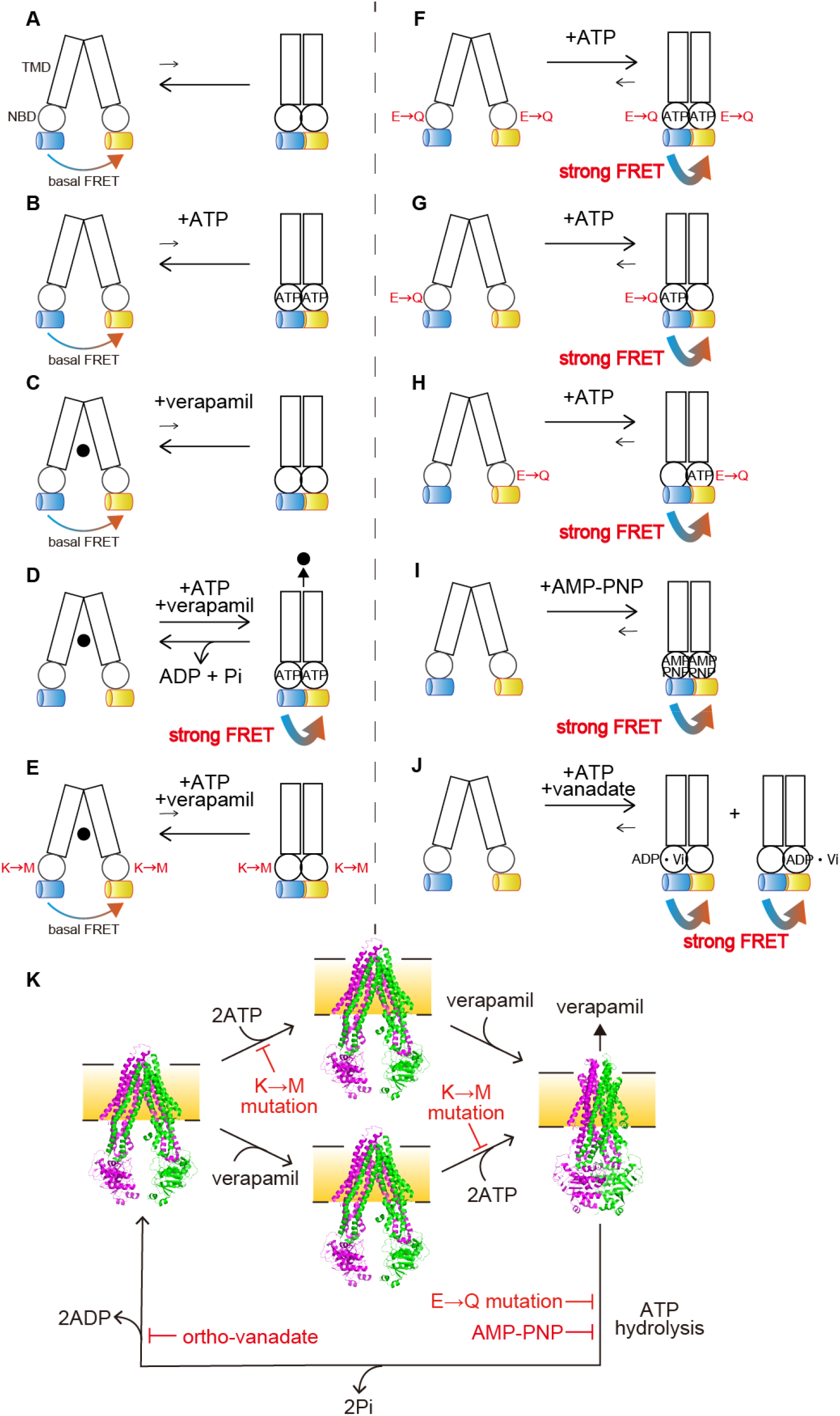
Schematic illustration of the ATP-dependent conformational changes of P-gp. (A-K) The transition equilibriums between the inward-facing conformation (left), which generates basal FRET, and the outward-facing conformation (right), which generates strong FRET, of human P-gp are indicated as arrows. Rectangles and circles represent TMDs and NBDs, respectively. Blue and yellow cylinders represent Cerulean and Venus, respectively. The replacement of Walker A lysine by methionine and of catalytic glutamate by glutamine are shown as M and Q, respectively. ●, verapamil. The inward-facing conformation is stable in the absence of ATP or verapamil (A, B, C). The presence of ATP and verapamil cause the transition from the inward-facing to the outward-facing conformation and generate strong FRET (D). Replacement of the Walker A lysine by methionine prevents ATP binding and generates basal FRET (E). Replacement of the Walker B glutamate by glutamine in either NBD stabilizes the outward-facing state and generates strong FRET (F, G, H). Binding of MgAMP-PNP stabilizes the outward-facing state and generate strong FRET (I). Orthovanadate traps P-gp in the stable post–ATP hydrolysis transition state (probably in either NBD) to generate strong FRET (J). (K) The model of transport cycle of P-gp caused by ATP. The inward-facing conformation is stable in the absence of ATP or verapamil (left and middle). The presence of ATP and verapamil cause the transition from the inward-facing to the outward-facing conformation (right). ATP hydrolysis and subsequent release of γ-phosphate from both NBDs allow P-gp to return to the inward-facing state. Mutations of the Walker A lysine prevents ATP binding. Mutations of the Walker B glutamate or binding of AMP-PNP inhibits ATP hydrolysis. Orthovanadate traps P-gp in the stable post–ATP hydrolysis transition state. The inward-facing conformation of mouse P-gp, (PDB: 4m1m); the outward-facing conformation of human P-gp, (PDB: 6c0v).

### ATP hydrolysis at both NBDs resets the conformation of P-gp to the low FRET state

Next, to investigate the role of ATP hydrolysis in the conformational change of P-gp, we analyzed P-gp(QQ)–FRET, which binds ATP but has impaired ATP hydrolysis. The proximity ratio of P-gp(QQ)–FRET was high in the presence of exogenously added ATP even without verapamil, but low in the absence of ATP (Fig. 3B). In addition, when ATP-depleted cells were incubated with the nonhydrolyzable ATP analogue, adenylyl-imidodiphosphate (AMP-PNP), the proximity ratio of P-gp–FRET was increased 1.5-fold even without verapamil treatment (Fig. 3C, 4I). It has been reported that P-gp is trapped in a state which mimics the post-hydrolysis transition state (18–20) when incubated with ATP and sodium orthovanadate (19, 20) and that the “vanadate trap” occurs at only one NBD randomly (19, 20). P-gp–FRET exhibited a high proximity ratio in the presence of ATP and sodium orthovanadate (Fig. 3C, 4J). Finally, we generated two single Walker B glutamate mutants, P-gp(QE)–FRET and P-gp(EQ)–FRET (Fig. 4G, H), in which the catalytic glutamate residue in only one NBD was substituted for glutamine. Both single Walker B mutants properly localized to the plasma membrane (Fig. S6A) but completely lost R6G transport activity (Fig. S6B). When these mutants were expressed in living cells, they exhibited a high proximity ratio even in the absence of a transport substrate (Fig. 2C). These results suggest that ATP hydrolysis and subsequent release of γ-phosphate from both NBDs allow the outward-facing state to return to the original inward-facing state.

## Discussion

Since the discovery of P-gp (9–11), numerous studies have been conducted to clarify the transport mechanism of P-gp. Most of these studies have utilized artificial membrane environments such as lipid-detergent mixed micelles (21), artificial lipid bilayers (5,7,22,23) or membrane vesicles derived from P-gp-overexpressing cells (24, 25). These conditions do not reflect physiological membrane environments in terms of lateral pressure, curvature, constituent lipid species, etc. Because these environmental factors can affect the function of P-gp, the detailed transport mechanism should be investigated in living cells. While a conformation sensitive monoclonal antibody, UIC2, which recognizes the extracellular loops of P-gp and specifically binds to the inward-facing structure (26), has been used to evaluate the conformational changes of the extracellular region of P-gp in living cells (6,27,28), an analysis of the conformational changes in living cells is technically difficult. In this study, we detected the change in distance between two NBDs of human P-gp in living cells by using FRET-based non-invasive imaging. Because the NBDs move dramatically in P-gp and the insertion of fluorescent proteins after the NBDs did not compromise the proper localization or substrate transport activity of P-gp, we expected the constructs to be suitable for monitoring the intrinsic behavior of P-gp in living cells.

P-gp–FRET on the plasma membrane in living cells showed a basal FRET signal (Fig. 1B, C). Considering that the ATP binding-deficient mutant P-gp(MM)–FRET showed a similar level of FRET as P-gp–FRET (Fig. 2A), ATP binding did not contribute to basal FRET. Alternatively, because the distance between the C-terminal of NBD1 and of NBD2 of human P-gp is about 30 Å in the inward-facing conformation (Movie_S01), it is possible that FRET occurs to some extent even in this conformation. The proximity ratio was increased by the addition of a transport substrate, verapamil, in a concentration-dependent manner, whereas the FRET signal of P-gp–VsCn was unchanged (Fig. 1C, D). The EC_50_ of the verapamil-stimulated proximity ratio in intact cells (3.7 ± 0.9 μM) (Figs. 1D) was comparable to the K_M_ (2.2 ± 0.2 – 4.1 ± 0.4 μM) of the verapamil-induced ATP hydrolysis by purified human P-gp (29). These results suggest that verapamil triggers the conformational change of P-gp and that Cerulean inserted after NBD1 and Venus fused after NBD2 approach each other in close proximity via dimerization of the two NBDs to increase intramolecular FRET (Fig. 4D).

To examine the role of ATP binding in the conformational change of P-gp, intracellular ATP was depleted from living cells by using streptococcal toxin streptolysin O (SLO). SLO is a member of the cholesterol-dependent pore-forming toxins. After binding to the plasma membrane, SLO oligomerizes and creates large β-barrel pores (17). In the presence of exogenously added ATP, the proximity ratio of P-gp–FRET was increased by the addition of verapamil, but not in the absence of ATP (Fig. 3A). These results suggest that both ATP and substrate binding are required for enhancing the conformational change from the inward-facing state (Fig. 4A-D). This hypothesis is consistent with the result of an ATP binding-deficient mutant P-gp(MM)–FRET in intact cells, which did not respond to verapamil (Fig. 2A). As with P-gp(QQ)–FRET, which we explain below, the results of wild-type and mutant P-gp are consistent with intact cells and SLO-treated ATP-depleted cells, suggesting that the ATP-dependent conformational change of P-gp is not affected by SLO treatment.

When ATP-depleted cells were incubated with the nonhydrolyzable ATP analogue AMP-PNP, the proximity ratio of P-gp–FRET was increased even without verapamil treatment (Fig. 3C). The same effect was seen with the ATPase-deficient mutant P-gp(QQ)–FRET in intact cells and ATP-depleted cells. This mutant’s proximity ratio was high in the presence of endogenous or exogenously added ATP independent of verapamil, but low in the absence of ATP (Fig. 3A, B). These results suggest that human P-gp moves spontaneously between the inward-facing and outward-facing structures at low frequency and that ATP before hydrolysis stabilizes the dimerized NBDs and fixes P-gp to the outward-facing structure (Fig. 4F, I). Furthermore, single Walker B glutamate mutants, P-gp(QE)–FRET and P-gp(EQ)–FRET, exhibited high proximity ratios even in the absence of a transport substrate (Fig. 2C), suggesting that inhibition of ATP hydrolysis at one NBD can stabilize dimerized NBDs and fixes P-gp to the outward-facing structure (Fig. 4G, H). This finding is consistent with biochemical analyses that showed orthovanadate traps P-gp in a state that mimics the post-hydrolysis transition state before the release of γ-phosphate (18–20) and that the “vanadate trap” occurs at only one NBD randomly (15,19,20,30). Accordingly, P-gp–FRET exhibited a high proximity ratio in the presence of ATP and sodium orthovanadate (Fig. 3C).

Our results are in agreement with cryo-EM (8), luminescence (7), and a variety of biochemical studies, in which mutations in a single ATP binding site intensely decreased the ATPase activity (6,31,32) and drug transport (33). Other studies also suggested that P-gp utilizes two ATP molecules in a single transport cycle (34, 35). By contrast, some studies reported that MgAMP-PNP does not cause conformational changes of P-gp (5, 21) and that ATP does not cause conformational changes of the Walker B glutamate mutant (21); these observations suggest that ATP hydrolysis causes a transition from the inward-facing to the outward-facing structure (5, 21). However, because those experiments were performed under artificial membrane environments, they might not reflect the behavior of P-gp in living cells.

To conclude, our results suggest that ATP hydrolysis and the subsequent release of γ-phosphate from both NBDs reset the conformation of P-gp from the outward-facing to the inward-facing state (Fig. 4K). Furthermore, our study underlines the necessity of experiments using living cells to understand the detailed mechanism of membrane proteins in addition to *in vitro* experiments.

## Experimental procedures

### Plasmid construction

All primer sequences are listed in Table 1. P-gp constructs were designed as MRP1 FRET constructs (36). pcDNA3.1P-gp was constructed by the insertion of cDNA encoding human P-gp in pcDNA3.1 (Thermo Fisher Scientific, Austin, TX, USA). To engineer P-gp–Venus, the pcDNA3.1(-)P-gp backbone and Venus cDNA were PCR-amplified using the primer pairs #1/#2 and #3/#4, respectively. To engineer P-gp–Cerulean, the pcDNA3.1(+)P-gp backbone, Cerulean cDNA and the following ten–amino acid linker (GS repeats) were PCR amplified using the primer pairs #5/#6 and #7/#8, respectively. The P-gp–FRET construct was generated by inserting Cerulean cDNA and the following ten–amino acid linker into pcDNA3.1(-)P-gp–Venus using the primer pairs #5/#6 and #7/#8, respectively. To engineer P-gp–VsCn, the pcDNA3.1(-)P-gp–Venus backbone and Cerulean cDNA were PCR-amplified using the primer pairs #1/#4 and #9/#10, respectively. The amplified fragments were ligated using an In-Fusion HD cloning kit (Clontech, Mountain View, CA, USA). pcDNA3.1(-)P-gp(MM; K433M/K1076M)-FRET, pcDNA3.1(-)P-gp(QQ; E556Q/E1201Q) and other mutants were generated using the corresponding pcDNA3.1(-)P-gp mutant backbones.

**Table 1.**
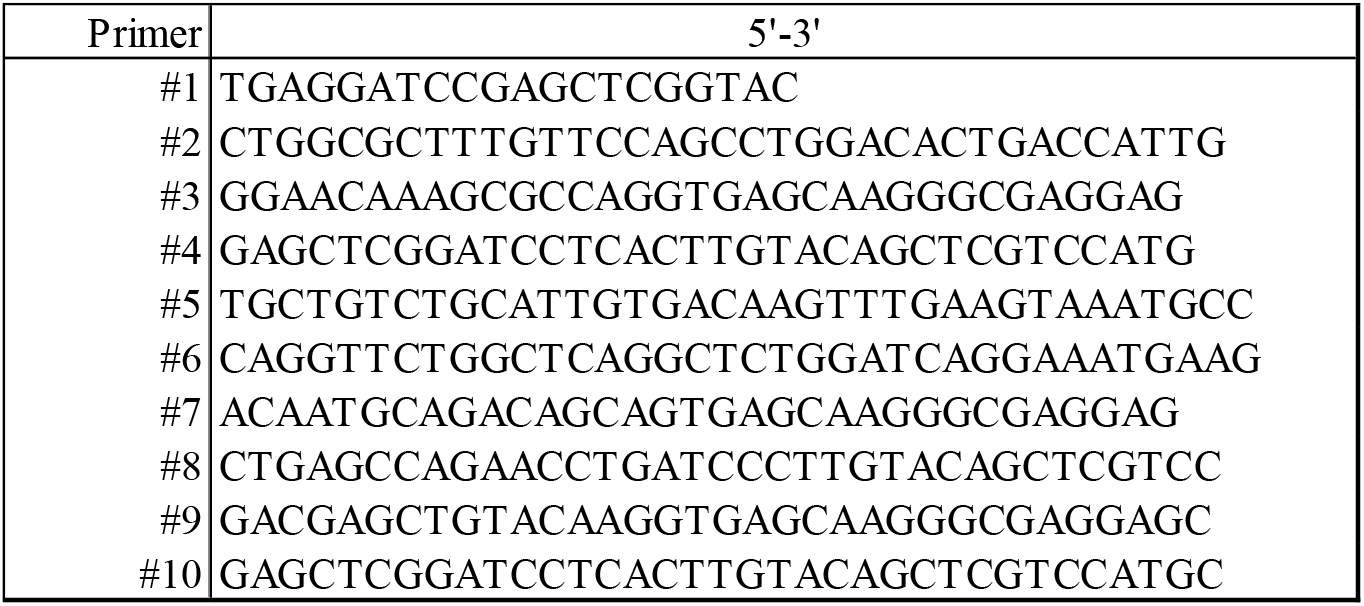
Primers

### Flow cytometry

HEK293 cells were cultured in Dulbecco’s modified Eagle’s medium (DMEM; Nacalai Tesque, Kyoto, Japan) supplemented with 10% fetal bovine serum (FBS; Gibco/Thermo) at 37°C under 5% CO_2_. HEK293 cells were seeded in 6-well plates (3.0 × 10^5^ cells/well) coated with poly-L-lysine (Sigma-Aldrich, St. Louis, MO, USA) and incubated for 24 h. P-gp constructs were transfected using Lipofectamine LTX Reagent (Thermo Fisher Scientific), and the transfected cells were incubated for an additional 24 h. Medium was replaced with serum-free DMEM containing 1 μM R6G with or without 25 μM PSC-833, a P-gp inhibitor (Abcam, Cambridge, MA, USA), and incubated for 30 min at 37°C under 5% CO_2_. Cells were washed twice with phosphate-buffered saline (PBS). After harvesting, the cells were washed with Hank’s balanced salt solution (HBSS; Gibco) and resuspended in HBSS. R6G accumulation was analyzed on an Accuri C6 flow cytometer (BD Biosciences, Franklin Lakes, NJ, USA).

### FRET microscopy

HEK293 cells were seeded at 3.0 × 10^5^ cells/well on glass-bottom dishes (AGC TECHNO GLASS, Shizuoka, Japan) coated with poly-L-lysine and incubated for 24 h. P-gp constructs were transfected using Lipofectamine LTX Reagent and incubated for an additional 23 h. Medium was then replaced by FluoroBrite DMEM (Gibco) supplemented with 10% FBS, sodium pyruvate (Gibco), and GlutaMAX (Gibco) and incubated for another 1 h. Cells were further incubated with 100 μM verapamil chloride (Fujifilm Wako Pure Chemical Corporation, Osaka, Japan) for 5 min at 37°C under 5% CO_2_ as necessary. Fluorescence images were acquired at 37°C under 5% CO_2_ by an LSM700 confocal microscope equipped with a Plan-Apochromat 63×/1.4 NA oil immersion objective lens (Carl Zeiss, Oberkochen, Germany). Cerulean was excited at 445 nm, and the fluorescence emission was collected using a short-pass filter (∼490 nm). Venus was excited at 488 nm, and the fluorescence emission was collected using a band-pass filter (521–600 nm). For the FRET signal, the laser was set to 445 nm, and the sensitized emission was collected using a band-pass filter (521–600 nm).

### Cell permeabilization

HEK293 cells were plated at 3.0 × 10^5^ cells/well on glass-bottom based dishes coated with poly-L-lysine and incubated for 24 h. P-gp constructs were transfected using Polyethyleneimine “Max” (MW: 40,000) (PEI-MAX; Polysciences, Warrington, PA, USA) and incubated for an additional 24 h. Cell permeabilization was performed as described previously (37) with slight modifications. Briefly, cells were washed twice with ice-cold PBS and incubated in serum-free DMEM containing 50 ng/μL SLO (BioAcademia, Osaka, Japan) for 5 min on ice, followed by three washes with ice-cold PBS. The cells were further incubated for 10 min at 37°C under 5% CO_2_ in transport buffer (25 mM HEPES-KOH [pH 7.4], 115 mM KOAc, 4.5 mM MgCl_2_) preincubated at 37°C containing 2 μg/mL DAPI (Sigma-Aldrich) in the presence or absence of 5 mM ATP disodium salt (Oriental Yeast, Tokyo, Japan). The cells were then washed twice with transport buffer and soaked in transport buffer pre-incubated at 37°C supplemented with 5 mM ATP disodium salt, 5 mM AMP-PNP lithium salt (Sigma-Aldrich), or 5 mM ATP disodium salt and 1 mM orthovanadate (Fujifilm Wako), as indicated. The cells were then incubated with 100 μM verapamil chloride for 5 min at 37°C under 5% CO_2_ as indicated. Only DAPI-positive cells used for the FRET microscopy. FRET microscopy was performed on an LSM700 microscope as described above.

### FRET calculation

FRET efficiency (proximity ratio) was calculated as follows:

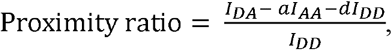

where *I*_DA_ is the FRET signal or the sensitized emission of the acceptor during donor excitation (Ex 445 nm/Em 521–600 nm), *I*_DD_ is the donor fluorescence during donor excitation (Ex 445 nm/Em ∼490 nm), and *I*_AA_ is the acceptor fluorescence during acceptor excitation (Ex 488 nm/Em 521–600 nm). a and d are the crosstalk coefficients of the acceptor and donor, respectively. These constants were calculated from P-gp–Venus and P-gp–Cerulean samples using the ImageJ plugin “FRET and Colocalization Analyzer”. The proximity ratio pseudocolour images were generated using MetaMorph software (Molecular Devices, San Jose, CA, USA). For quantification of the proximity ratio, the plasma membrane was determined from automatically segmented Venus images of P-gp–Venus, P-gp–FRET, and P-gp–VsCn or Cerulean images of P-gp–Cerulean using ImageJ-based Fiji software. The proximity ratio was quantified on a cell-by-cell basis.

### Protein expression and purification

Human P-gp cDNA was inserted into the expression vector pcDNA3.1(-). The C-terminus of the DNA was modified by elimination of the natural termination codon and insertion of a *Not*I restriction cleavage site, TEV protease cleavage site (ENLYFQG), ten histidine codons, a six–amino acid linker (GAAGTS), and the FLAG tag (DYKDDDDK), followed by a termination codon. FreeStyle 293-F cells (Thermo) were transfected with the resultant P-gp construct using PEI-MAX. After 48 h of transfection, the cells were harvested and resuspended in buffer A (50 mM HEPES-Na [pH 7.2], 150 mM NaCl, 50 mM KCl, 20% glycerol) containing 0.5 μM 2-mercaptethanol (Fujifilm Wako) and protease inhibitor (Roche, Basel, Switzerland) and solubilized with 0.6% (w/v) *n*-dodecyl-*β*-D-maltopyranoside (DDM; Anatrace, Maumee, OH). Solubilized proteins were applied to FLAG-M2 agarose (Sigma-Aldrich) pre-equilibrated with buffer A. The mixture was rotated for 18 h. The resin was washed with a 4× bed volume of buffer A containing 0.6% DDM three times. The resin was further washed with a 4× bed volume of buffer A containing 0.1% DDM three times. The protein was eluted from the resin with 1× bed volume of buffer A containing 0.1% DDM, 300 μg/mL FLAG peptide (Sigma-Aldrich), and 3× FLAG peptide (Sigma-Aldrich). The eluate was concentrated by centrifugation using an Amicon Ultra (Merck, Kenilworth, NJ, USA). Purified protein was stored at −80°C. All purification steps were performed at 0–4°C.

### Reconstitution into proteoliposomes and measurement of ATPase activity

Purified protein was reconstituted in lipid as described previously(38) with some modifications. Briefly, L-α-lecithin from egg yolk (Fujifilm Wako) was resuspended in reaction buffer (50 mM Tris-HCl (pH7.4), 0.1 mM EGTA) at a final concentration of 20 mg/mL. The suspension was sonicated in a bath sonicator. The purified protein (2.1 μg) was mixed with 40 μg lipid, and the mixture was incubated on ice for 20 min. Proteoliposomes were incubated at 37°C for 30 min with 3 mM ATP disodium salt and 5 mM MgCl_2_ in 20 μL reaction buffer. Verapamil chloride, DAPI, and BeFx (10 mM BeSO_4_, 50 mM NaF) were added to the reaction buffer at 100 μM, 2 μg/mL, and 1 mM, respectively. ATPase reaction was stopped by the addition of 20 μL of 20 mM EDTA. The ATPase activity was estimated by measuring the amount of released ADP by HPLC using a titanium column(39).

### Statistical analysis

Data are presented as means ± S.E. Multiple comparisons were evaluated using Tukey’s post hoc test following one-way ANOVA. All statistical analyses were performed using Origin2018 software (LightStone, Tokyo, Japan)

## Acknowledgements

This work was supported by JSPS KAKENHI Grant Number 18H05269 to KU and 18K19176 to YK.

## Supporting Information Legends

**Movie_S01**

The animation illustrates the overall structural change of P-gp between the inward-(PBD code 4Q9H) and outward-facing (PBD code 6C0V) structures. Structures are aligned based on NBD1 structures. The linear distance between Cα atoms of the C-terminal residues of NBD1 and NBD2 are shown as a yellow dashed line. The morphing movie was calculated using the ‘morph’ command of PyMOL 1.8.

**Fig. S1.**
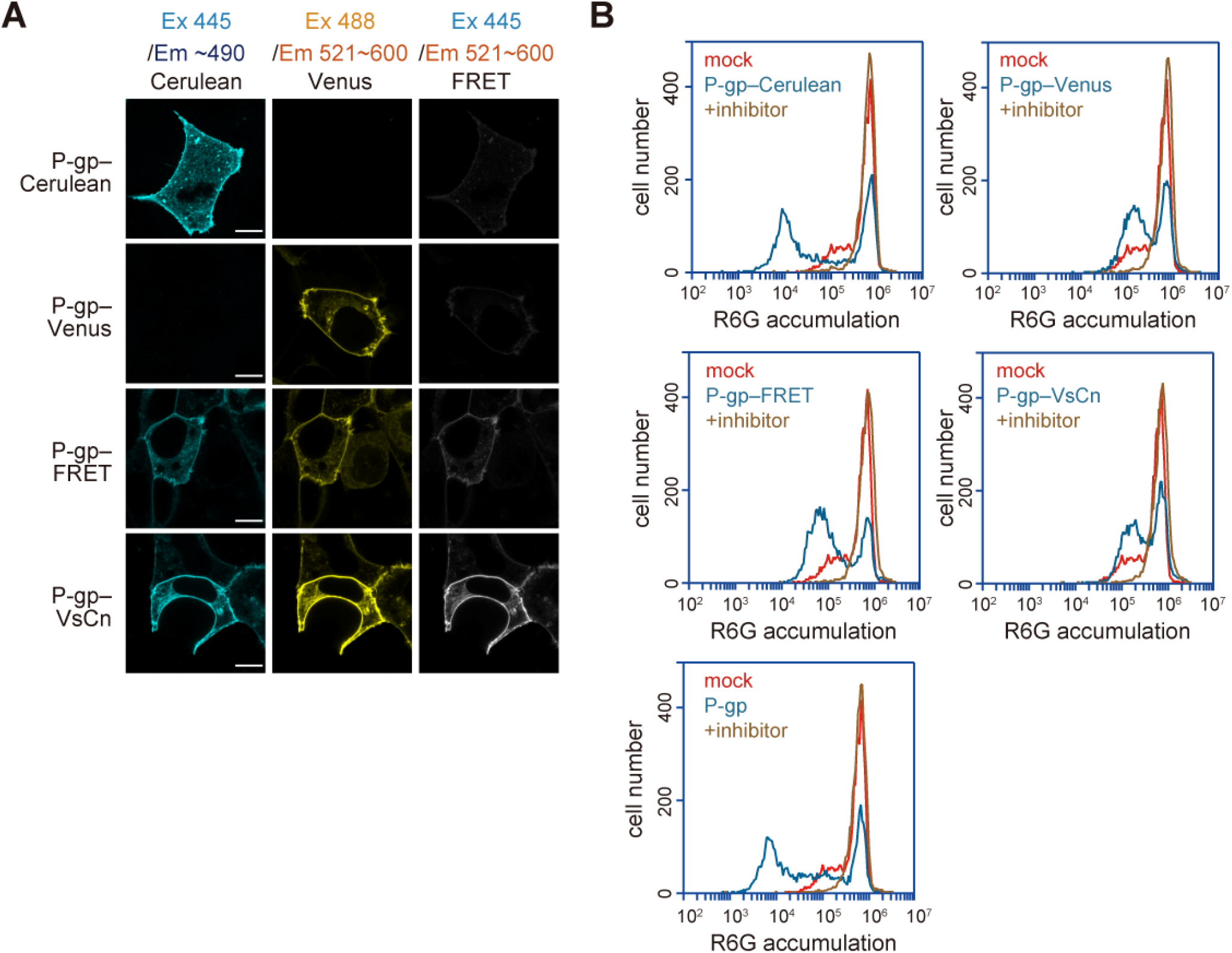
The insertion and fusion of fluorescent proteins does not impair the localization or function of P-gp. (**A**) Fluorescent protein–tagged P-gps were transiently expressed in HEK293 cells. Fluorescent images were obtained at 37°C using an LSM700 confocal microscope (Zeiss). Scale bars, 10 μm. (**B**) HEK293 cells transiently expressing fluorescent protein–tagged P-gps were incubated with 1 μM R6G with or without 25 μM inhibitor (PSC-833) for 30 min at 37°C. R6G accumulation in the cells was measured by flow cytometry.

**Fig. S2.**
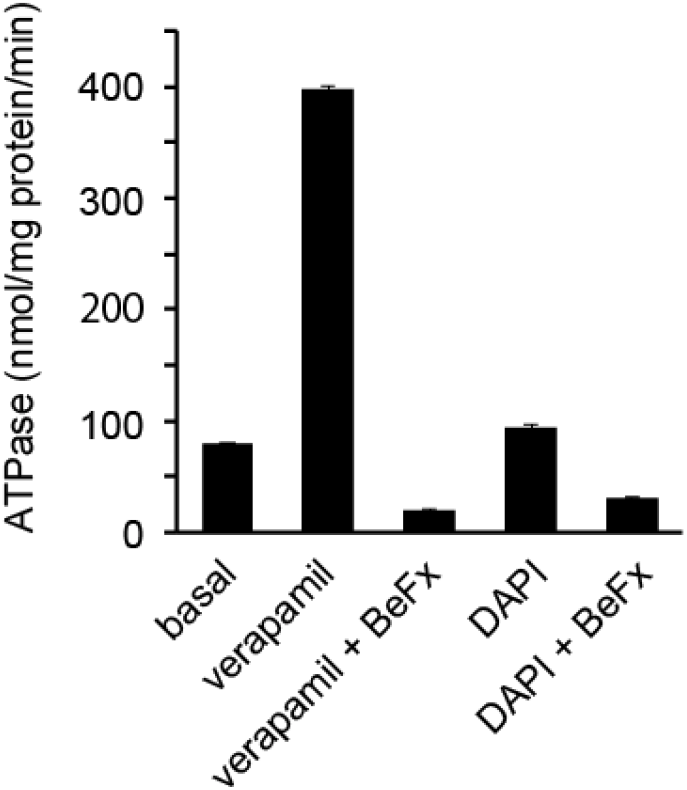
Verapamil, a transport substrate, stimulated the ATPase activity of P-gp, but DAPI, which is not a transport substrate, did not. Purified human P-gp was reconstituted in liposomes consisting of L-α-lecithin from egg yolk. Proteoliposomes were incubated for 30 min at 37°C with 3 mM ATP, 5 mM Mg^2+^, 100 µM verapamil chloride, 2 µg/mL DAPI, and 1 mM beryllium fluoride (BeFx, an ATPase inhibitor). ATPase activity was estimated by measuring the amount of released ADP. Data are expressed as means ± S.E.

**Fig. S3.**
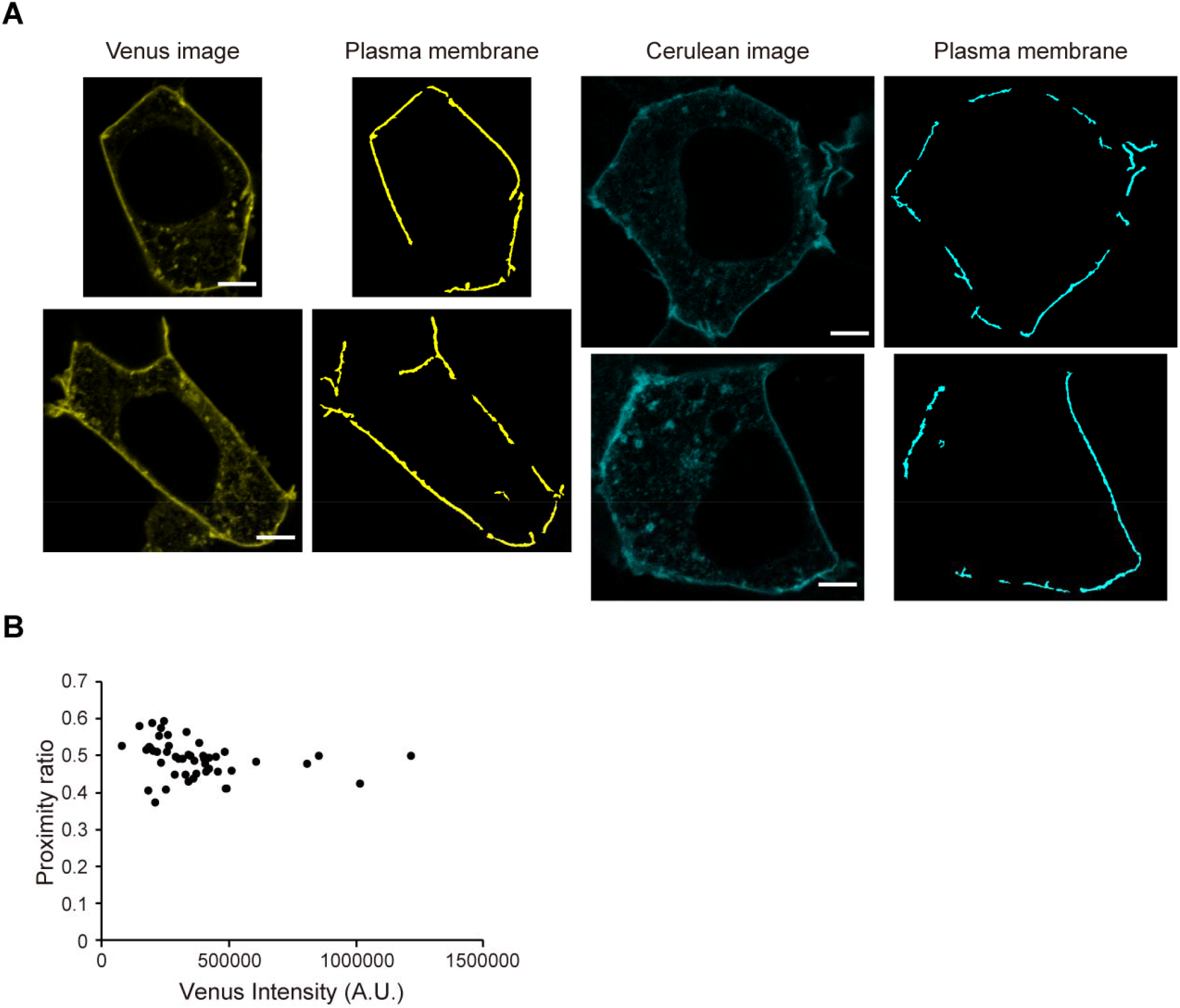
(**A**) The region of the plasma membrane determined from automatically segmented Venus images of P-gp–FRET and Cerulean images of P-gp–Cerulean. Scale bars, 5 µm. (**B**) The protein expression level had no significant effect on the proximity ratio of P-gp–FRET.

**Fig. S4.**
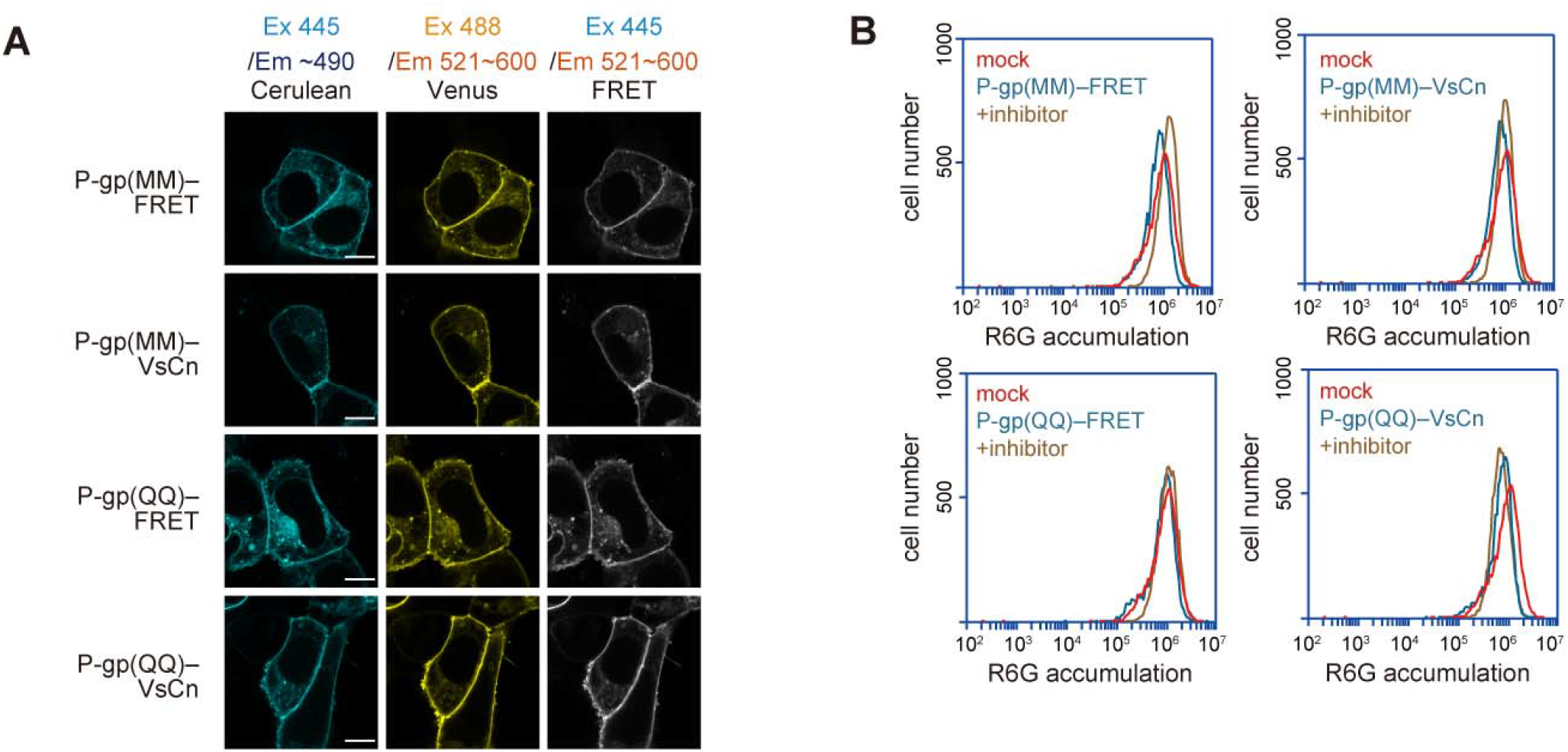
(**A**) Subcellular localization and FRET of P-gp(MM)–FRET and P-gp(QQ)–FRET. (**B**) R6G transport activity. Scale bars, 10 μm.

**Fig. S5.**
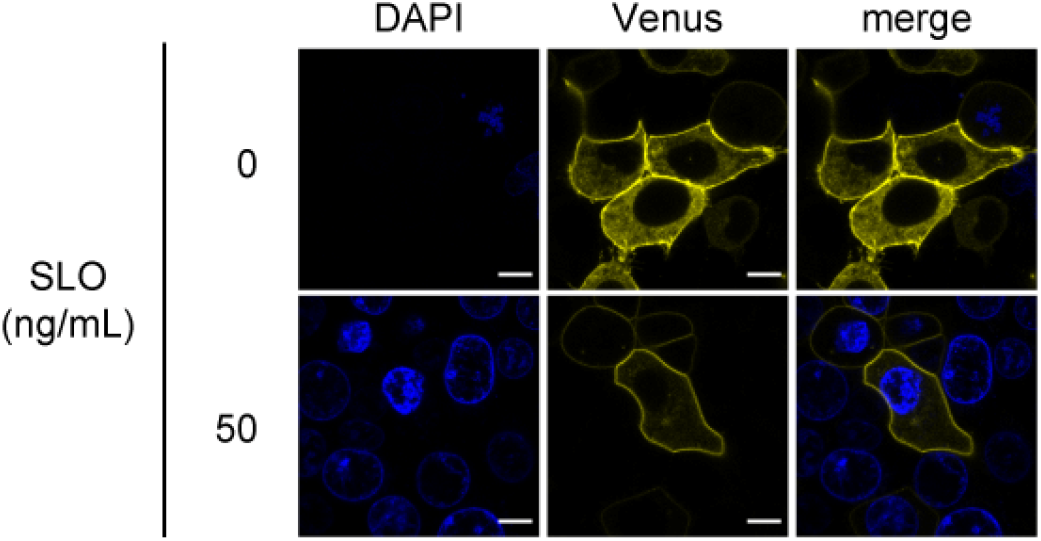
Subcellular localization of P-gp–Venus and DAPI staining in cells permeabilized with 50 ng/mL SLO. Fluorescent images were obtained at 37°C on a Nikon C2 confocal system equipped with a Plan Apochromat 40×/ dry objective lens. Scale bars, 10 μm.

**Fig. S6.**
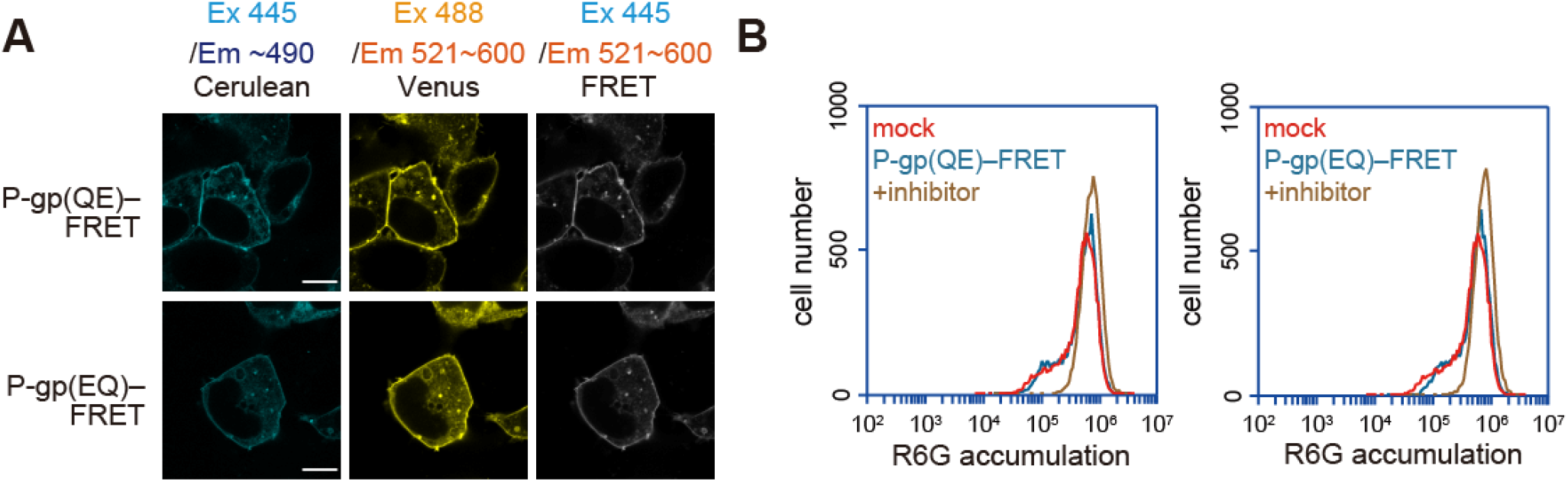
(**A**) Subcellular localization and FRET of P-gp(QE)–FRET and P-gp(EQ)–FRET. (**B**) R6G transport activity. Scale bars, 10 μm.

## References

1. Ueda, K., Cardarelli, C., Gottesman, M. M., and Pastan, I. (1987) Expression of a full-length cDNA for the human “MDR1” gene confers resistance to colchicine, doxorubicin, and vinblastine. Proc Natl Acad Sci U S A 84, 3004–3008

2. Gottesman, M. M., and Ling, V. (2006) The molecular basis of multidrug resistance in cancer: the early years of P-glycoprotein research. FEBS Lett 580, 998–1009

3. Schinkel, A. H., Smit, J. J., van Tellingen, O., Beijnen, J. H., Wagenaar, E., van Deemter, L., Mol, C. A., van der Valk, M. A., Robanus-Maandag, E. C., and te Riele, H. P. (1994) Disruption of the mouse mdr1a P-glycoprotein gene leads to a deficiency in the blood-brain barrier and to increased sensitivity to drugs. Cell 77, 491–502

4. Li, J., Jaimes, K. F., and Aller, S. G. (2014) Refined structures of mouse P-glycoprotein. Protein Sci 23, 34–46

5. Moeller, A., Lee, S. C., Tao, H., Speir, J. A., Chang, G., Urbatsch, I. L., Potter, C. S., Carragher, B., and Zhang, Q. (2015) Distinct conformational spectrum of homologous multidrug ABC transporters. Structure 23, 450–460

6. Bársony, O., Szalóki, G., Türk, D., Tarapcsák, S., Gutay-Tóth, Z., Bacsó, Z., Holb, I. J., Székvölgyi, L., Szabó, G., Csanády, L., Szakács, G., and Goda, K. (2016) A single active catalytic site is sufficient to promote transport in P-glycoprotein. Sci Rep 6, 24810

7. Zoghbi, M. E., Mok, L., Swartz, D. J., Singh, A., Fendley, G. A., Urbatsch, I. L., and Altenberg, G. A. (2017) Substrate-induced conformational changes in the nucleotide-binding domains of lipid bilayer-associated P-glycoprotein during ATP hydrolysis. J Biol Chem 292, 20412–20424

8. Kim, Y., and Chen, J. (2018) Molecular structure of human P-glycoprotein in the ATP-bound, outward-facing conformation. Science

9. Juliano, R. L., and Ling, V. (1976) A surface glycoprotein modulating drug permeability in Chinese hamster ovary cell mutants. Biochim Biophys Acta 455, 152–162

10. Chen, C. J., Chin, J. E., Ueda, K., Clark, D. P., Pastan, I., Gottesman, M. M., and Roninson, I. B. (1986) Internal duplication and homology with bacterial transport proteins in the mdr1 (P-glycoprotein) gene from multidrug-resistant human cells. Cell 47, 381–389

11. Ueda, K., Cornwell, M. M., Gottesman, M. M., Pastan, I., Roninson, I. B., Ling, V., and Riordan, J. R. (1986) The mdr1 gene, responsible for multidrug-resistance, codes for P-glycoprotein. Biochem Biophys Res Commun 141, 956–962

12. Kodan, A., Yamaguchi, T., Nakatsu, T., Sakiyama, K., Hipolito, C. J., Fujioka, A., Hirokane, R., Ikeguchi, K., Watanabe, B., Hiratake, J., Kimura, Y., Suga, H., Ueda, K., and Kato, H. (2014) Structural basis for gating mechanisms of a eukaryotic P-glycoprotein homolog. Proc Natl Acad Sci U S A 111, 4049–4054

13. Kodan, A., Yamaguchi, T., Nakatsu, T., Matsuoka, K., Kimura, Y., Ueda, K., and Kato, H. (2019) Inward- and outward-facing X-ray crystal structures of homodimeric P-glycoprotein CmABCB1. Nat Commun 10, 88

14. Kimura, Y., Matsuo, M., Takahashi, K., Saeki, T., Kioka, N., Amachi, T., and Ueda, K. (2004) ATP hydrolysis-dependent multidrug efflux transporter: MDR1/P-glycoprotein. Curr Drug Metab 5, 1–10

15. Szabó, K., Welker, E., Bakos Müller, M., Roninson, I., Váradi, A., and Sarkadi, B. (1998) Drug-stimulated nucleotide trapping in the human multidrug transporter MDR1. Cooperation of the nucleotide binding domains. J Biol Chem 273, 10132–10138

16. Tombline, G., Bartholomew, L. A., Urbatsch, I. L., and Senior, A. E. (2004) Combined mutation of catalytic glutamate residues in the two nucleotide binding domains of P-glycoprotein generates a conformation that binds ATP and ADP tightly. J Biol Chem 279, 31212–31220

17. Stewart, S. E., D’Angelo, M. E., Paintavigna, S., Tabor, R. F., Martin, L. L., and Bird, P. I. (2015) Assembly of streptolysin O pores assessed by quartz crystal microbalance and atomic force microscopy provides evidence for the formation of anchored but incomplete oligomers. Biochim Biophys Acta 1848, 115–126

18. Hofmann, S., Januliene, D., Mehdipour, A. R., Thomas, C., Stefan, E., Brüchert, S., Kuhn, B. T., Geertsma, E. R., Hummer, G., Tampé, R., and Moeller, A. (2019) Conformation space of a heterodimeric ABC exporter under turnover conditions. Nature 571, 580–583

19. Urbatsch, I. L., Sankaran, B., Weber, J., and Senior, A. E. (1995) P-glycoprotein is stably inhibited by vanadate-induced trapping of nucleotide at a single catalytic site. J Biol Chem 270, 19383–19390

20. Urbatsch, I. L., Tyndall, G. A., Tombline, G., and Senior, A. E. (2003) P-glycoprotein catalytic mechanism: studies of the ADP-vanadate inhibited state. J Biol Chem 278, 23171–23179

21. Verhalen, B., Dastvan, R., Thangapandian, S., Peskova, Y., Koteiche, H. A., Nakamoto, R. K., Tajkhorshid, E., and Mchaourab, H. S. (2017) Energy transduction and alternating access of the mammalian ABC transporter P-glycoprotein. Nature 543, 738–741

22. Verhalen, B., Ernst, S., Börsch, M., and Wilkens, S. (2012) Dynamic ligand-induced conformational rearrangements in P-glycoprotein as probed by fluorescence resonance energy transfer spectroscopy. J Biol Chem 287, 1112–1127

23. Dastvan, R., Mishra, S., Peskova, Y. B., Nakamoto, R. K., and Mchaourab, H. S. (2019) Mechanism of allosteric modulation of P-glycoprotein by transport substrates and inhibitors. Science 364, 689–692

24. Liu, R., and Sharom, F. J. (1996) Site-directed fluorescence labeling of P-glycoprotein on cysteine residues in the nucleotide binding domains. Biochemistry 35, 11865–11873

25. Qu, Q., and Sharom, F. J. (2001) FRET analysis indicates that the two ATPase active sites of the P-glycoprotein multidrug transporter are closely associated. Biochemistry 40, 1413–1422

26. Alam, A., Küng, R., Kowal, J., McLeod, R. A., Tremp, N., Broude, E. V., Roninson, I. B., Stahlberg, H., and Locher, K. P. (2018) Structure of a zosuquidar and UIC2-bound human-mouse chimeric ABCB1. Proc Natl Acad Sci U S A 115, E1973–E1982

27. Mechetner, E. B., Schott, B., Morse, B. S., Stein, W. D., Druley, T., Davis, K. A., Tsuruo, T., and Roninson, I. B. (1997) P-glycoprotein function involves conformational transitions detectable by differential immunoreactivity. Proc Natl Acad Sci U S A 94, 12908–12913

28. Druley, T. E., Stein, W. D., and Roninson, I. B. (2001) Analysis of MDR1 P-glycoprotein conformational changes in permeabilized cells using differential immunoreactivity. Biochemistry 40, 4312–4322

29. Kimura, Y., Kioka, N., Kato, H., Matsuo, M., and Ueda, K. (2007) Modulation of drug-stimulated ATPase activity of human MDR1/P-glycoprotein by cholesterol. Biochem J 401, 597–605

30. Urbatsch, I. L., Beaudet, L., Carrier, I., and Gros, P. (1998) Mutations in either nucleotide-binding site of P-glycoprotein (Mdr3) prevent vanadate trapping of nucleotide at both sites. Biochemistry 37, 4592–4602

31. Müller, M., Bakos, E., Welker, E., Váradi, A., Germann, U. A., Gottesman, M. M., Morse, B. S., Roninson, I. B., and Sarkadi, B. (1996) Altered drug-stimulated ATPase activity in mutants of the human multidrug resistance protein. J Biol Chem 271, 1877–1883

32. Loo, T. W., and Clarke, D. M. (1995) Covalent modification of human P-glycoprotein mutants containing a single cysteine in either nucleotide-binding fold abolishes drug-stimulated ATPase activity. J Biol Chem 270, 22957–22961

33. Azzaria, M., Schurr, E., and Gros, P. (1989) Discrete mutations introduced in the predicted nucleotide-binding sites of the mdr1 gene abolish its ability to confer multidrug resistance. Mol Cell Biol 9, 5289–5297

34. Eytan, G. D., Regev, R., and Assaraf, Y. G. (1996) Functional reconstitution of P-glycoprotein reveals an apparent near stoichiometric drug transport to ATP hydrolysis. J Biol Chem 271, 3172–3178

35. Shapiro, A. B., and Ling, V. (1997) Positively cooperative sites for drug transport by P-glycoprotein with distinct drug specificities. Eur J Biochem 250, 130–137

36. Iram, S. H., Gruber, S. J., Raguimova, O. N., Thomas, D. D., and Robia, S. L. (2015) ATP-Binding Cassette Transporter Structure Changes Detected by Intramolecular Fluorescence Energy Transfer for High-Throughput Screening. Mol Pharmacol 88, 84–94

37. Ogasawara, F., Kano, F., Murata, M., Kimura, Y., Kioka, N., and Ueda, K. (2019) Changes in the asymmetric distribution of cholesterol in the plasma membrane influence streptolysin O pore formation. Sci Rep 9, 4548

38. Ishigami, M., Tominaga, Y., Nagao, K., Kimura, Y., Matsuo, M., Kioka, N., and Ueda, K. (2013) ATPase activity of nucleotide binding domains of human MDR3 in the context of MDR1. Biochim Biophys Acta 1831, 683–690

39. Kimura, Y., Shibasaki, S., Morisato, K., Ishizuka, N., Minakuchi, H., Nakanishi, K., Matsuo, M., Amachi, T., Ueda, M., and Ueda, K. (2004) Microanalysis for MDR1 ATPase by high-performance liquid chromatography with a titanium dioxide column. Anal Biochem 326, 262–266

